# Adversarial random forests for omics synthesis

**DOI:** 10.64898/2026.09.09.750490

**Authors:** Cesaire J. K. Fouodo, Jan Kapar, Anke Hüls, Donghai Liang, Marvin N. Wright

## Abstract

Data availability is critical for understanding complex disease pathways and developing robust predictive models. Although high-throughput omics technologies have improved insight into disease mechanisms, data acquisition from inaccessible tissues such as the central nervous system remains a major limitation, causing small sample sizes and complicating early prediction of neurodegenerative disorders such as Alzheimer’s and Parkinson’s diseases. Generative modeling has emerged as a powerful approach for synthesizing data to support downstream clustering and prediction with small sample size, but existing methods rarely handle high-dimensional tabular omics data effectively. Adversarial random forests (ARFs) provide a well-performing framework for tabular data generation but are not designed for high-dimensional settings. To address this limitation, we introduce high-dimensional ARF (*h*-ARF), an extension of ARF optimized for integrated clinical and high-dimensional omics data. Using benchmarks across nine datasets and eight performance metrics, we show that *h*-ARF better preserves both feature distributions, and downstream clustering and prediction utilities compared with ARFs. The method is implemented in the opensource R package harf, available on CRAN.

## 1 Introduction

Over the last decade, the usage of omics data, including DNA methylation, transcriptomic, metabolomic, and proteomic data, has become increasingly important for understanding the biological mechanisms underlying complex diseases. Although the collection of omics data has become less expensive (1, 2), a major limitation for numerous diseases remains the accessibility of the organs or tissues primarily affected. Typical examples are neurodegenerative diseases such as Alzheimer’s disease (AD) and Parkinson’s disease (PD), which are characterized by progressive neuronal damage and complex metabolic dysregulation (3, 4). For such diseases, collecting omics data from brain tissue is preferred. However, because brain tissue is not readily accessible for living patients, omics data for AD and PD are typically derived from post-mortem samples, resulting in limited sample sizes and reduced utility for early disease prediction and crosstissue biomarker discovery. Furthermore, because complex disease mechanisms involve environmental exposures as well as demographic, anthropometric, and clinical covariates, integrating these multi-modal inputs with omics data presents a substantial challenge due to their heterogeneous joint distributions and varied feature dimensionalities.

Generative modeling has recently emerged as an effective technique for addressing sample-size limitations in statistical modeling (5, 6). Its fundamental principle leverages empirical patterns within a dataset to synthesize novel data with similar characteristics. For example, this technique has been shown to be effective for improving prediction performance using clinical data of breast cancer and diabetes patients (7). Classical approaches based on parametric assumptions, such as the multivariate normal distribution, often struggle to capture complex dependencies in omics data, particularly in the presence of mixed variable types (continuous and categorical) and non-linear joint structures. More flexible approaches, based on deep learning and probabilistic modeling, including variational autoencoders (VAEs), generative adversarial networks (GANs), diffusion models, and transformer-based architectures, have therefore been proposed (8–13). However, these methods are typically designed for image, text, or signal data and are data-intensive, limiting their effectiveness in small-sample tabular omics settings. Similarly, alternative probabilistic frameworks such as Bayesian networks rely on accurate estimation of high-dimensional dependencies and often require substantial sample sizes to perform reliably (14). Methods such as TabSyn (13) extend tabular data generation using latent diffusion with variational autoencoders to model inter-feature dependencies in a continuous latent space. However, like other deep learning-based approaches, they remain computationally intensive and require large sample sizes.

Adversarial random forests (ARFs) have recently been introduced as a non-parametric approach for density estimation and generative modeling based on random forests (15, 16). They are well suited for tabular data and have shown strong performance in small samples. ARFs support conditional sampling, enabling flexible data generation under feature constraints.

Despite their attractiveness, ARFs remain largely restricted to low-to-moderate-dimensional settings, with benchmark studies typically involving only a few dozen features (< 60) (15, 17, 18). In contrast, omics data are commonly highdimensional and exhibit complex correlation structures that are not well captured in such settings, limiting the applicability of ARFs for realistic biomedical data generation. While pre-filtering or variable selection may be used to reduce dimensionality, these approaches risk discarding weak but jointly informative signals, particularly in polygenic contexts where effects are small and distributed across many features. This limitation is further amplified in multimodal data integration, where high-dimensional omics features may dominate learning at the expense of more informative low-dimensional clinical variables (19–21). Consequently, effective generative modeling in this setting requires methods that explicitly account for differences in modality dimensionality.

We propose the *h*-ARF algorithm, an extension of ARF for high-dimensional integrative generative modeling. We present its theoretical formulation, provide a software package, and evaluate its performance on benchmark datasets for clustering and prediction, as well as in an Alzheimer’s disease (AD) case study.

## 2 Methods

We consider a dataset **D** = ( **X**_omx_, **X**_clin_ ), where **X**_omx_ *∈* ℝ^*n*×*p*^ represents the omics data and **X**_clin_ the ( *n* × *q*) -dimensional clinical data. We assume that the omics modality is potentially high-dimensional, with *p ≫ n* quantitative features, whereas the clinical modality is low-dimensional, with *q ≪ n* features that may be mixed (categorical and continuous). Our objective is to generate a synthetic dataset 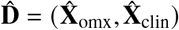 that shares similar joint structure with the original dataset **D**. To this end, we aim to train a generative model that approximates the joint distribution ℙ (**D**) and generates synthetic observations from the learned distribution 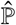.

### 2.1 Adversarial random forests (ARFs)

The key ideas of adversarial random forests (ARFs) build on unsupervised random forests (URFs) (22) and generative adversarial networks (GANs), combining random forest-based partitioning with a discriminator–generator framework (23). URFs extend RFs to an unsupervised setting by creating an artificial classification problem that distinguishes original from synthetic data. ARFs proceed as follows:

1. Based on the URF framework, construct an artificial supervised learning problem, by generating a naive synthetic dataset 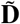 from the marginal distribution of the original feature space. Combine the original and synthetic datasets to form 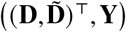 *∈* ℝ ^2*n*×( *p*+*q*+1)^, where **Y** is a binary indicator distinguishing real from synthetic observations. This extended dataset is used to train a supervised RF model 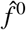 to predict **Y**.
2. If 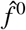, the current discriminator, is not able to discriminate real from synthetic observations given a predefined predicting performance measure (e.g., an accuracy of 0.5 + *δ* (*δ* > 0) (24)) and a maximum number of iterations, generate new synthetic data by sampling from leaves of generator 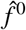 (generator). Subsequently, train a new 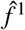 to distinguish between the original and the current synthetic data.
3. Repeat updating the generator and the discriminator until either a maximal number of iteration is reached or the prediction accuracy of the RF model 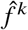 drops down to or below 0.5 + *δ*; meaning the ARF algorithm converged.
4. Forde (Forest Density Estimation): Estimate the joint probability distribution *p*^*l*^ within each leaf under the assumption that, via the recursive partitioning of the original dataset, features are marginally distributed within leaves.
5. Forge (Forest data Generation): Randomly draw a decision tree and a leaf *l*, weighted by the sample size of the original observations that ended up in *l*. Subsequently, sample an observation from the estimated joint probability distribution *p*^*l*^ within the leaf *l*.

Convergence is crucial for the ARF algorithm because synthetic observations are generated under the assumption that features are locally independent within terminal nodes. As the discriminator’s ability to distinguish real from synthetic observations decreases, the synthetic distribution approaches the original joint distribution (ℙ (**D**) ). This implies that the recursive partitioning induced by the forest has produced leaves in which the local independence assumption is sufficiently satisfied. This assumption can be more challenging to be satisfied in high- than in low-dimensional settings. In low-dimensional settings, observations are more densely distributed, increasing the likelihood that correlated features are jointly independent within leaves after partioning. In high-dimensional settings, the sample size is often insufficient to adequately explore the joint feature space. For example, (a) if different feature subsets drive similar correlation patterns, the RF classifier may randomly ignore some subsets during training, (b) even in case all subsets may be used, the sample size might often be insufficient to decompose the entire joint correlation structure and obtain features that are jointly independent within leaves. As a result, correlated features are less likely to be jointly independent within leaves as the dimensionality increases, violating the independence assumption of the ARF algorithm. Consequently, the Forde stage may fail to adequately approximate joint distribution within leaves.

Beyond the convergence challenges encountered in highdimensional settings, integrating omics (high-dimensional) and clinical (low-dimensional) data may pose another obstacle to ARF to learn the joint distribution ℙ (**D** )adequately. The high dimensionality of omics features can dominate the learning process, potentially masking the structure driven by clinical variables. As a result, clinically relevant variables may be underutilized, even when they are more strongly associated with the response variable of interest (21).

### 2.2 High-dimensional ARFs

The high-dimensional ARF (*h*-ARF) algorithm is an extension of the ARF framework tailored to high-dimensional settings. The objective of *h*-ARF is to improve the applicability of ARF in high-dimensional data by increasing the likelihood that the local independence assumptions underlying ARF are satisfied within terminal nodes. Our framework relies on partitioning the omics feature space. Although omics datasets often exhibit complex correlation structures, these dependencies are typically not uniformly distributed across all features. Instead, they are largely driven by underlying biological and genetic architectures, such as co-regulated genes, metabolic pathways, or shared molecular functions. Consequently, omics features frequently form groups with stronger within-group than between-group dependencies. This structure motivates partitioning the high-dimensional feature space into lower-dimensional chunks, within which the local independence assumptions underlying ARF are more likely to hold. The remaining dependencies across chunks are subsequently captured through latent representations. To simplify the idea behind the *h*-ARF algorithm, consider two random variables *A* and *C*. We also consider a random variable *Z* such that *A ⊥ C* | *Z*, which yields the following factorization of the joint distribution

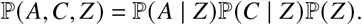

Therefore sampling *Z ∼* ℙ (*Z*) followed by conditional sampling *A* | *Z ∼* ℙ (*A* | *Z*) and *C* | *Z ∼* ℙ (*C* | *Z*) reconstructs the original joint distribution ℙ (*A, C*) . Hence, the variable *Z* can be interpreted as an auxiliary representation capturing the dependence structure between *A* and *C*. Ideally, the auxiliary variable *Z* should allow the factorization of ℙ (*A, C, Z*) . In practice, *Z* could be approximated by any representation that captures the shared dependence structure between *A* and *C*. Since ARF enables conditional sampling, one of the milestones of *h*-ARF is to construct such a variable *Z*.

To extend the idea above to high-dimensional settings, we decompose the original feature space into low-dimensional subspaces in which the ARF algorithm can converge reliably. We consider a function Ψ – any clustering algorithm, for instance – that partitions the omics space into *R* homogeneous isolated regions. We deliberately avoid using disjoint terminology here, as these regions may share common distributional structures. We further assume the existence of a function Φ so that

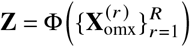

is a set of meta-features summarizing the shared structure across regions in a low-dimensional latent space. Details about Ψ and Φ are presented later. We rewrite the original joint distribution as

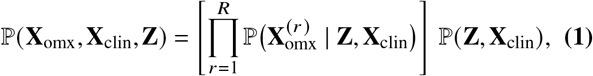

where 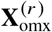 denotes the subset of omics features beloging to the isolated region *r ∈* {1, …, *R*} . Algorithm 1 outlines the framework for fitting a *h*-ARF model. For each region *r*, given latent and clinical features, the ARF algorithm is used to learn the underlying region-specific distribution. We denote by ( *f* ^*r*^, *p*^*r*^ ), the coupl e Forge and Forde for the regionspecific distribution 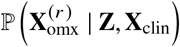, and by ( *f* ^*c*^, *p*^*c*^), the fitted Forge and Forde for ℙ (**Z, X**_clin_ ). Trained ARF models are subsequently used in algorithm 2 to generate new observations, by first sampling 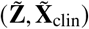, followed 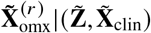 from each isolated region.

#### Algorithm 1

*h*-ARF generators and density estimators

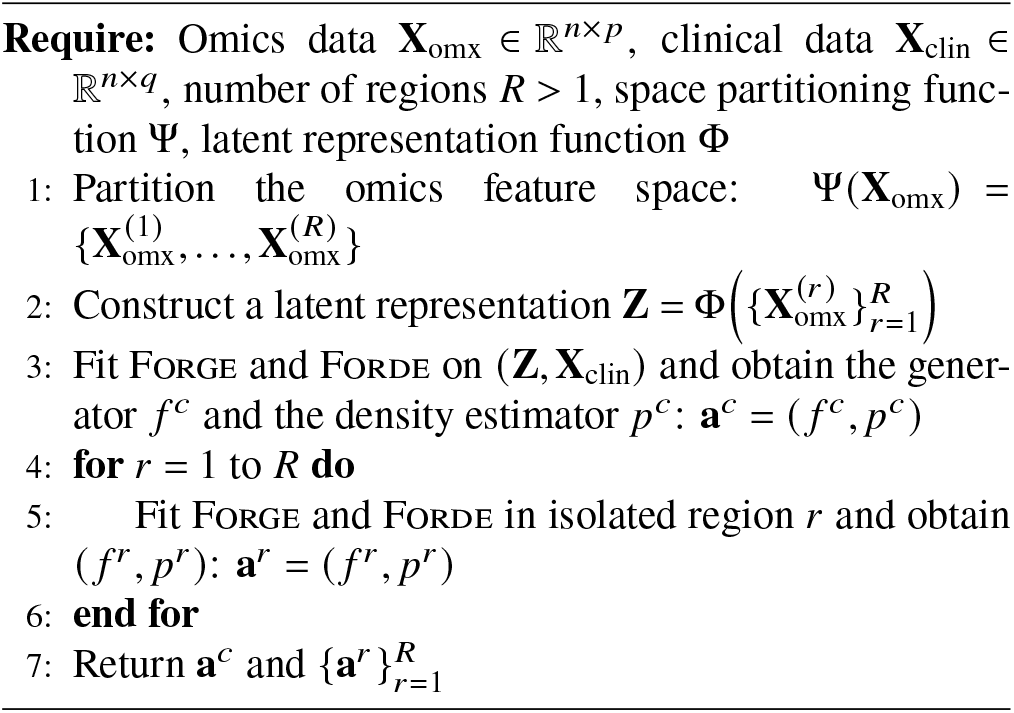

#### Algorithm 2

*h*-ARF data synthesizer

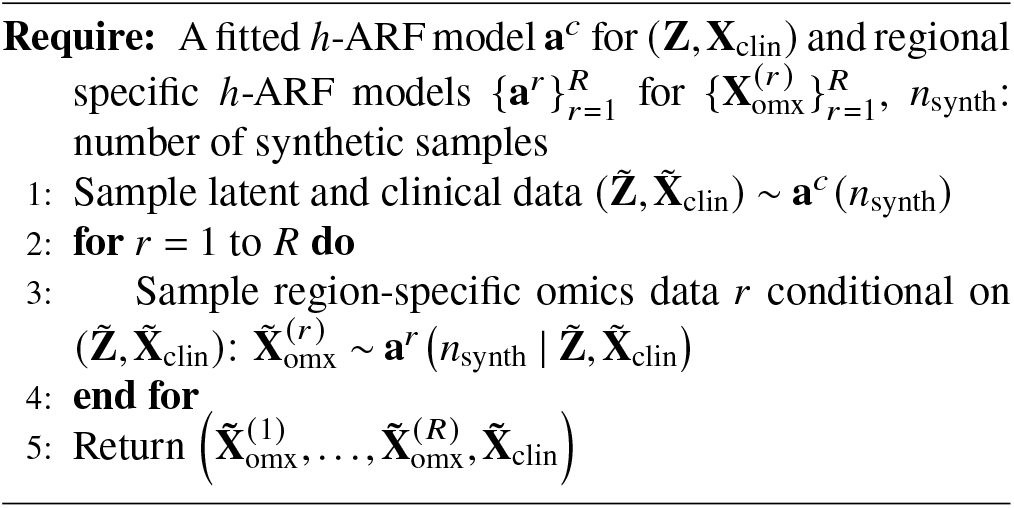

#### 2.2.1 Feature partitioning via Ψ

We explain how we partition the feature space via the function Ψ. This procedure is inspired by the previous works of (25). Starting from the transposed omics matrix 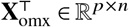, we compute a Spearman similarity matrix between features.

We perform principal component analysis (PCA) via singular value decomposition (SVD) to obtain a low-dimensional representation of the space of the omics features. The number of principal components is selected using the elbow method (26). In the reduced space, omics features are projected and clustered into *k* disjoint groups using the *k*-means algorithm, which yields an initial partition of the feature space. We emphasize that the choice of *k* does not guarantee that each region is sufficiently low-dimensional for ARF convergence. Therefore, assuming the resulting groups are sufficiently homogeneous, regions with dimensionality exceeding a predefined chunk size are further subdivided into smaller subsets.

#### 2.2.2 Construction of the latent space via Φ

The meta-features **Z** are designed to capture the joint dependency structure across the isolated regions. Our procedure for finding the joint space is based on canonical correlation analysis (CCA) (27, 28), which aims to identify block components that summarize the relevant information both between and within blocks of features. We distinguish between unsupervised and supervised settings in the construction of **Z**. For the unsupervised setting, the goal is to find unit norm block weight vectors solving

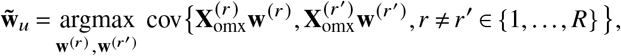

with 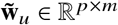 the basis vectors of the latent space and *m* the number of components of interest, **w**^(*r* )^an element vector with so many columns as the dimension of 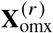 . The metafeatures are given by 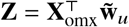. Whereas CCA captures correlations between data blocks without considering the response block, partial least squares (PLS) performs supervised integration by incorporating the response variables into the latent component estimation. The PLS algorithm achieves this by solving

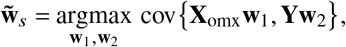

**Y** being the set of response variables, 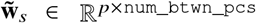, **w**_1_ *∈* R^*p*×1^, and **w**_2_ a column vector with as many elements as the dimension of **Y**.

There are numeruous approaches to solve the CCA optimization problem. We utilize the regularized approach proposed by (28) and implemented in the `R` package `RGCCA` (version 3.0.3) (29). We use the implementation proposed in the R package `pls` (version 2.9-0) to solve the PLS problem in a supervised setting (30).

## 3 Benchmark study with downstream clustering and prediction datasets

We evaluate the proposed framework with ARF as baseline method in two application settings: a downstream clustering evaluation using publicly available reference cell-type datasets, and a downstream prediction benchmark study based on publicly available gene expression data with simulated outcomes. Clustering groups cells into distinct, homogeneous types, whereas prediction trains models to infer the outcome. Computation was conducted on a single CPU node of the Emory University High-Performance Computing (HPC) cluster, running Rocky Linux (version 8) with R version 4.3. For each study, runs were parallelized using the R package `batchtools,` version 0.9.18 (31).

### 3.1 Downstream clustering with benchmark datasets

Four single-cell gene expression datasets are used for clustering analysis (Table 1), with cell type treated as non omics variable. Data preprocessing follows (25), including removal of genes with zero expression across all cells or expressed in fewer than 6% of cells. The genes were further filtered to retain the top 10% of those highly expressed.

**Table 1.**
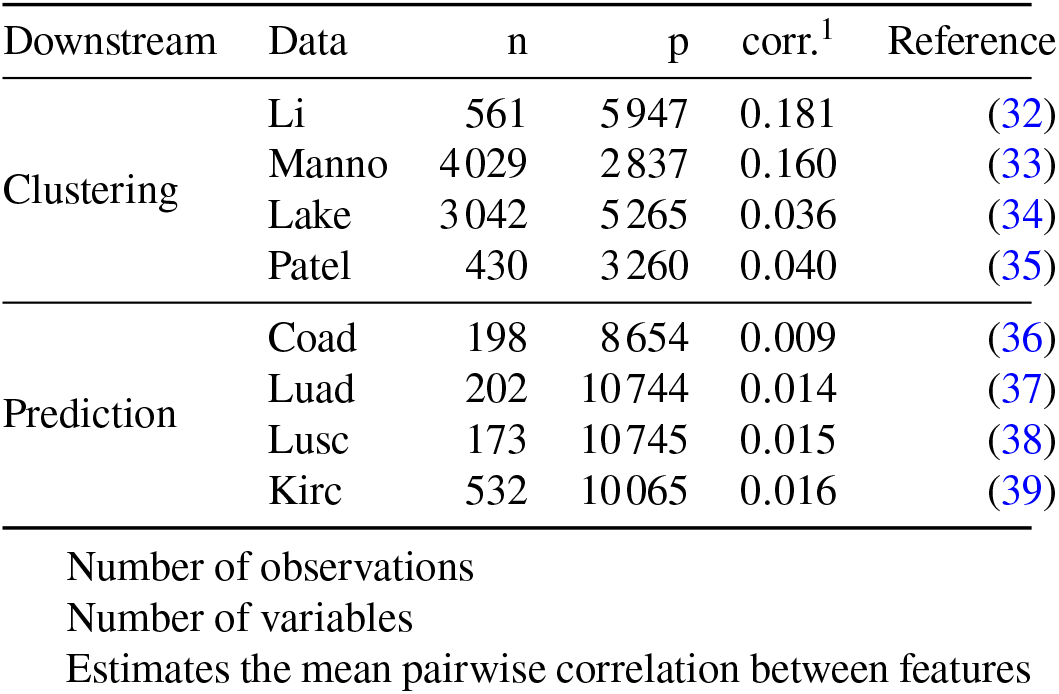
Summary description of datasets.

| Downstream | Data | n | p | corr.<sup>1</sup> | Reference |
| --- | --- | --- | --- | --- | --- |
| Clustering | Li | 561 | 5947 | 0.181 | (32) |
|  | Manno | 4029 | 2837 | 0.160 | (33) |
|  | Lake | 3042 | 5265 | 0.036 | (34) |
|  | Patel | 430 | 3260 | 0.040 | (35) |
| Prediction | Coad | 198 | 8654 | 0.009 | (36) |
|  | Luad | 202 | 10744 | 0.014 | (37) |
|  | Lusc | 173 | 10745 | 0.015 | (38) |
|  | Kirc | 532 | 10065 | 0.016 | (39) |

Number of observations
Number of variables
Estimates the mean pairwise correlation between features

We assess ARF and *h*-ARF on downstream clustering datasets over 100 runs. In each run, we use a 70:30 split for training and testing sets, and synthesize another training data of same size as the original training set. Performance is evaluated using six measures grouped into distributional similarity, clustering utility, and runtime. Distributional measures include univariate Wasserstein distance (UVD) (40), maximum mean discrepancy (MMD) with radial basis kernel, and pairwise Pearson correlation distance (CD), where UVD quantifies average feature-wise Wasserstein distance between original and synthetic data, MMD measures discrepancies in a reproducing kernel Hilbert space, and CD captures pairwise correlation structure differences. Clustering utility is assessed using adjusted Rand index (ARI) (41) and normalized mutual information (NMI) (42), which quantify agreement between clustering structures induced by synthetic and original data. To estimate these performance measures, we run the *k*-means clustering algorithm on the test cells. Subsequently, we use the training and synthetic datasets to fit two RF classification models, with cell types as response variables. Hyperparameters, including the number of omics feature groups and chunk size, were selected independently of the test data to avoid potential data leakage. Using the fitted RF models, we predict the cell types of the testing samples and estimate the ARI and NMI between the clustered cell types using *k*-means and the predicted cell types. We compute the difference between the ARI obtained from the original data and that obtained from the synthetic data, as well as the NMI. Finally, we report the runtime for synthesizing data using each algorithm. For this experimental setup, ARF and *h*-ARF are implemented with an ensemble size of 10 decision trees. To analyze the impact of chunk size, the *h*-ARF parameter varies from 5 to 50 in increments of 5.

### 3.2 Downstream prediction with benchmark datasets

We use four datasets from The Cancer Genome Atlas (TCGA) (36–39, 43) in the downstream prediction analysis. These include colon adenocarcinoma (Coad), lung adenocarcinoma (Luad), lung squamous cell carcinoma (Lusc), and kidney renal clear cell carcinoma (Kirc), covering major cancer types across tissues. Coad, Luad, and Lusc focus on transcriptomic signatures of tumor subtypes, while Kirc represents one of the most common and aggressive kidney cancer forms. Clinical covariates include age and sex, along with a dichotomized artificial response constructed from a logistic model, since original tumor stages were highly imbalanced. Specifically, we simulate the response using a logit model with effects drawn from *N* (0, 0.5) assigned to 0.01% of randomly selected omics features and age.

For downstream prediction, we quantify the synthetic data using UVD, MMD, and CD, as well as the area under the ROC curve (AUC) using RF and Lasso (44) as final classifiers. We trained both RF and Lasso on the original and the synthesized datasets to mitigate potential model leakage stemming from our RF-base generative frameworks. We also assess the distribution similarity of the synthetized datasets using UVD, MMD, and CD. Evaluation is conducted across 100 simulations. In each run, the baseline dataset is divided into 70:30 training and testing sets. Predictive models are subsequently developed using the training subset. We synthesize training data using the ARF and the *h*-ARF algorithm. Subsequently, we predict the response variable using the trained models and estimate the difference in AUC between the two predictive models. The closer the expected difference is to zero, the better the utility of downstream predictive analysis on the synthetic training datasets.

## 4 Results of benchmark studies

Results are presented for downstream clustering and prediction for publicly available datasets.

### 4.1 Downstream clustering

Figure 1 shows the estimated performance measures for clustering-designed datasets for a chunk size of 5 in *h*-ARF. The *h*-ARF algorithm outperforms the ARF algorithm in downstream clustering datasets. Across all datasets, *h*-ARF provides better data quality for UVD, MMD and CD. Regarding downstream data utility performance measures, the ARI difference between the original and synthetic data is consistently larger for ARF across all datasets, indicating the main difference between the two algorithms. The synthetic Li and Manno datasets – with more strongly correlated features – showed better performance than the synthetic Lake and Patel datasets for *h*-ARF and ARF. The substantial improvements in both distributional similarity and downstream utility clustering performance come at the cost of increased computational runtime, highlighting a strong accuracy–efficiency trade-off for *h*-ARF (see table 2).

**Table 2.** Summary of runtime in minutes across clustering and prediction datasets for a chunk size of 5 for *h*-ARF.

| Clustering datasets |  |  | Prediction datasets |  |  |
| --- | --- | --- | --- | --- | --- |
| Dataset | *h*-ARF | ARF | Dataset | *h*-ARF | ARF |
| Li | 9.54 | 1.02 | Coad | 5.30 | 0.03 |
| Manno | 9.50 | 0.90 | Luad | 11.72 | 0.08 |
| Lake | 15.46 | 2.50 | Lusc | 28.78 | 0.10 |
| Patel | 8.50 | 0.83 | Kirc | 2.90 | 0.01 |

**Fig 1.**
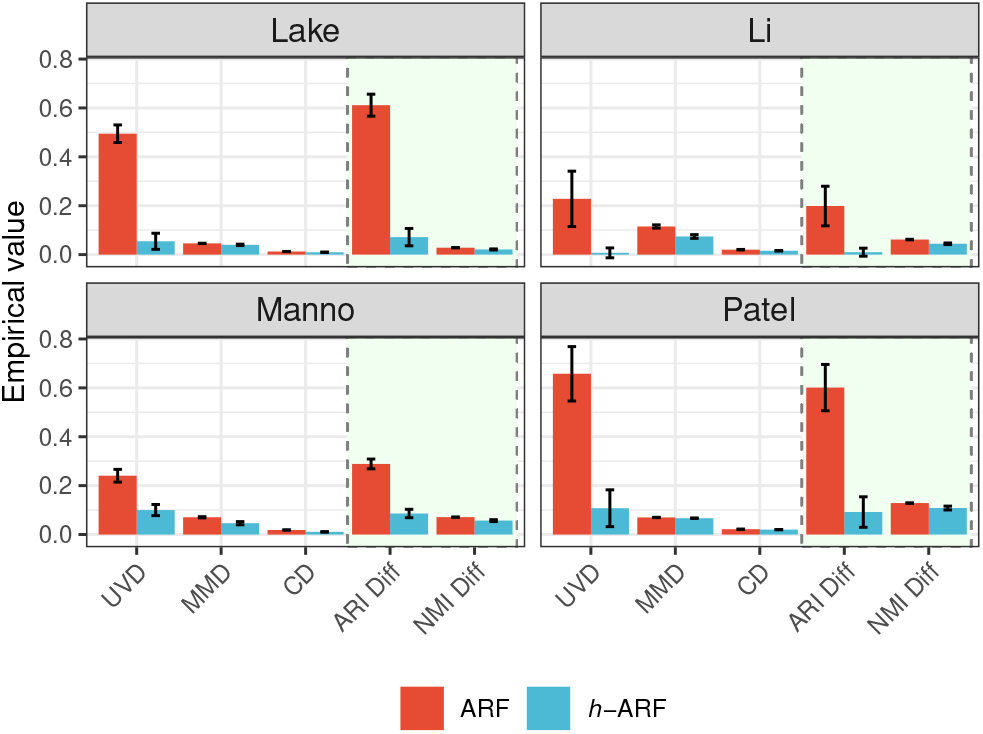
Estimated performance measures across the four clustering-designed datasets for the *h*-ARF (chunk size of 5) and ARF algorithms. Distributional similarity measures are shown on the left (white background), whereas downstream clustering performance measures are shown on the right (light cyan background). Values closer to zero indicate better performance.

Results illustrating the effect of varying chunk size on the *h*- ARF algorithm are provided in the supplementary materials (figure 5). This effect is consistent across all datasets: increasing the chunk size reduces the quality of synthetic data, in both distributional fidelity and downstream utility, although reducing the runtime (figure 6). We further observe that this sensitivity depends on the data’s underlying correlation structure, with weakly correlated datasets being less affected than strongly correlated ones.

### 4.2 Downstream prediction

The *h*-ARF method consistently outperforms ARF in downstream prediction (Figure 2). Across all four datasets, AUC differences are more tightly centered around zero for *h*-ARF, indicating better preservation of predictive signal. This trend holds for both RF and Lasso models, whereas ARF-generated data leads to substantial performance degradation, particularly in the Kirc dataset, which exhibits stronger feature correlations. Distribution-based metrics show no substantial differences between methods (Figure 3).

**Fig 2.**
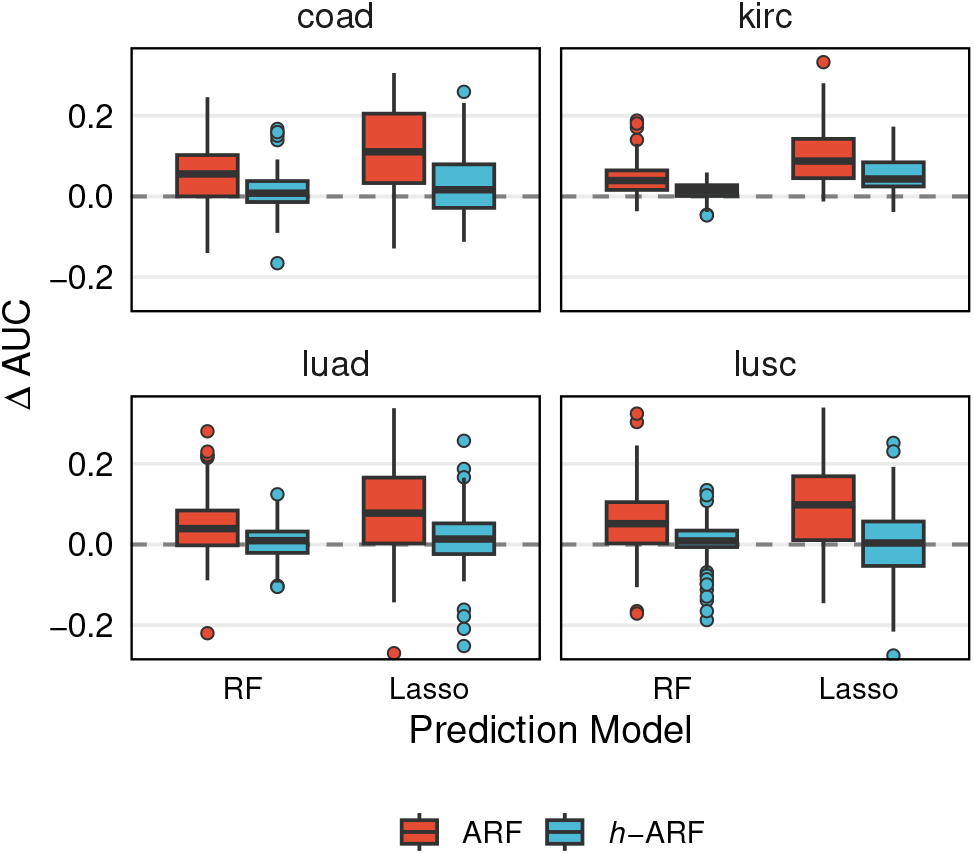
Performance in downstream prediction analysis showing differences in AUC between models trained on original versus synthetic data (RF and Lasso). The *h*- ARF uses chunk size 5, and distributions centered closer to zero indicate better predictive utility preservation.

**Fig 3.**
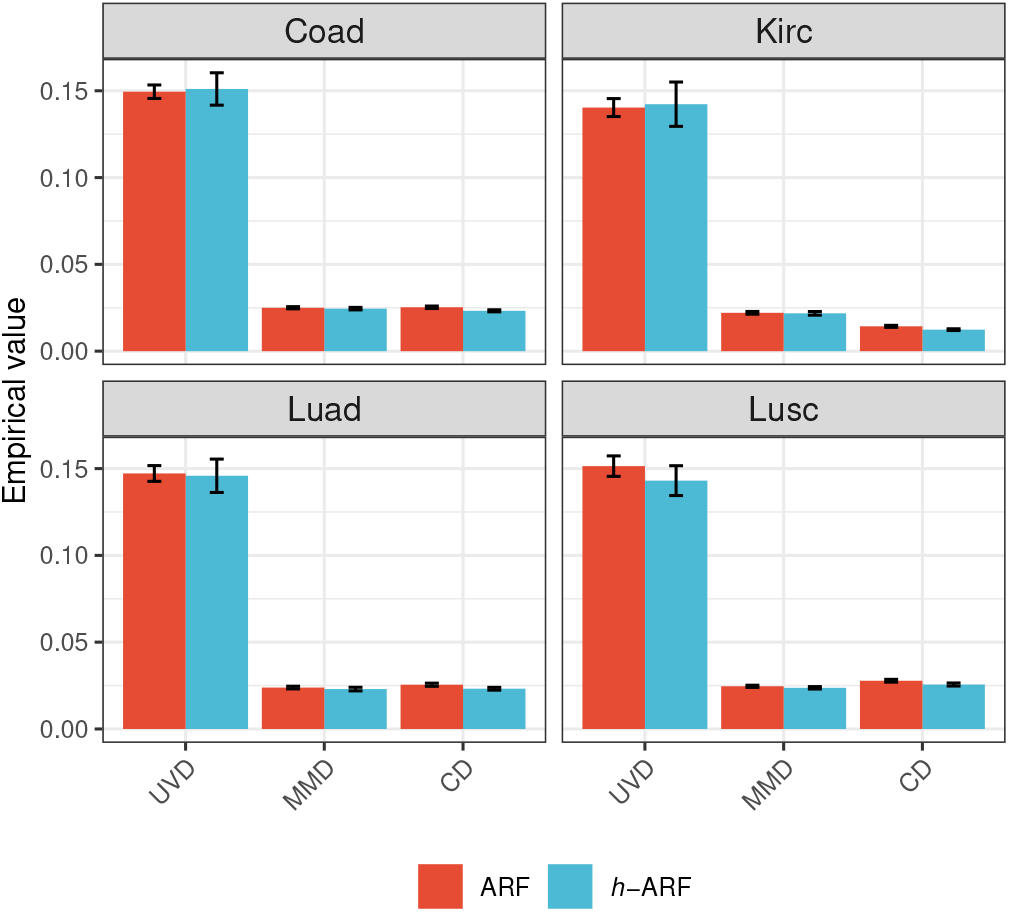
Estimated distribution performance measures for ARF and *h*-ARF for the downstream prediction datasets.

**Fig 4.**
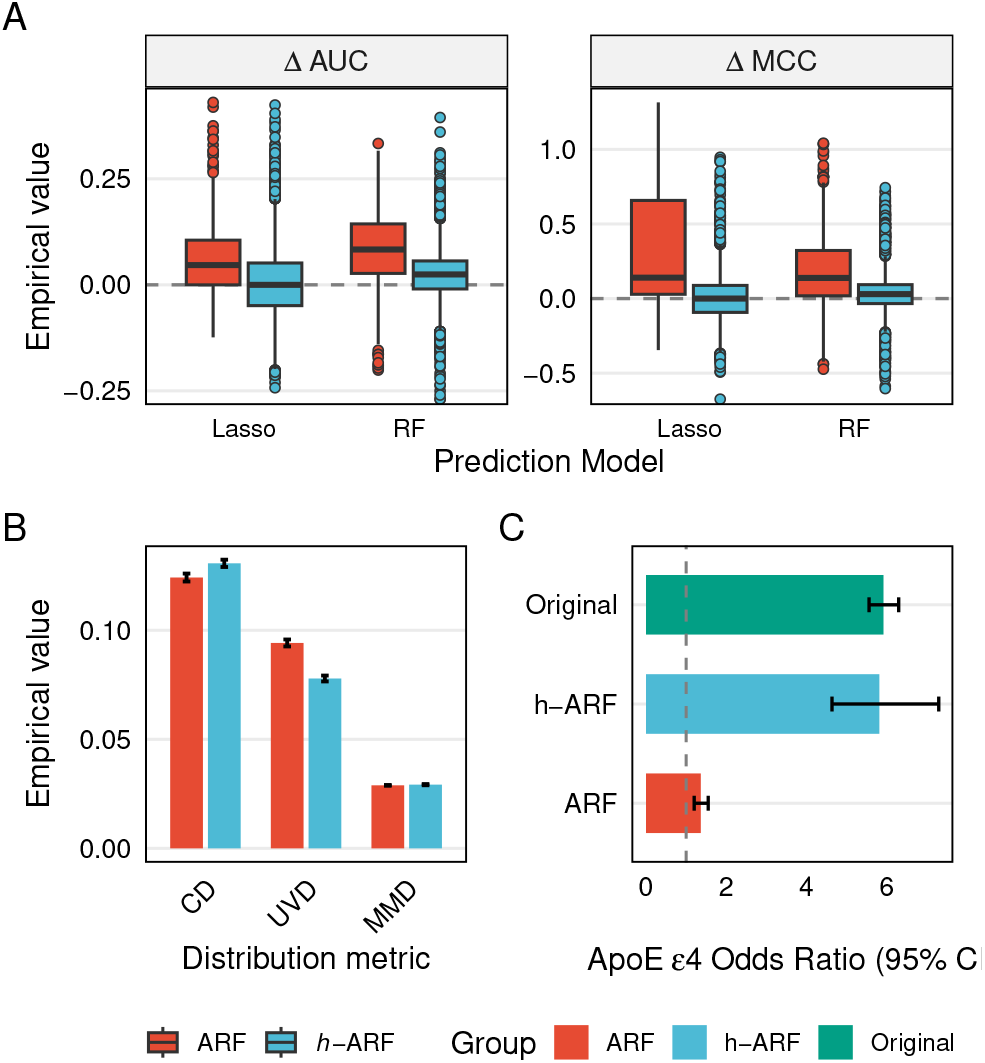
Synthetic data quality for the AD metabolomics dataset evaluated through downstream predictive analyses. (A) Differences in MCC and AUC between models trained on original versus synthetic datasets are shown for both Lasso and RF classifiers. (B) Empirical distribution-based performance metrics are presented to assess preservation of feature distributions. (C) Estimated odds ratios for (*ApoE* ) are shown for models fitted on the original dataset and on the synthesized datasets.

**Fig 5.**
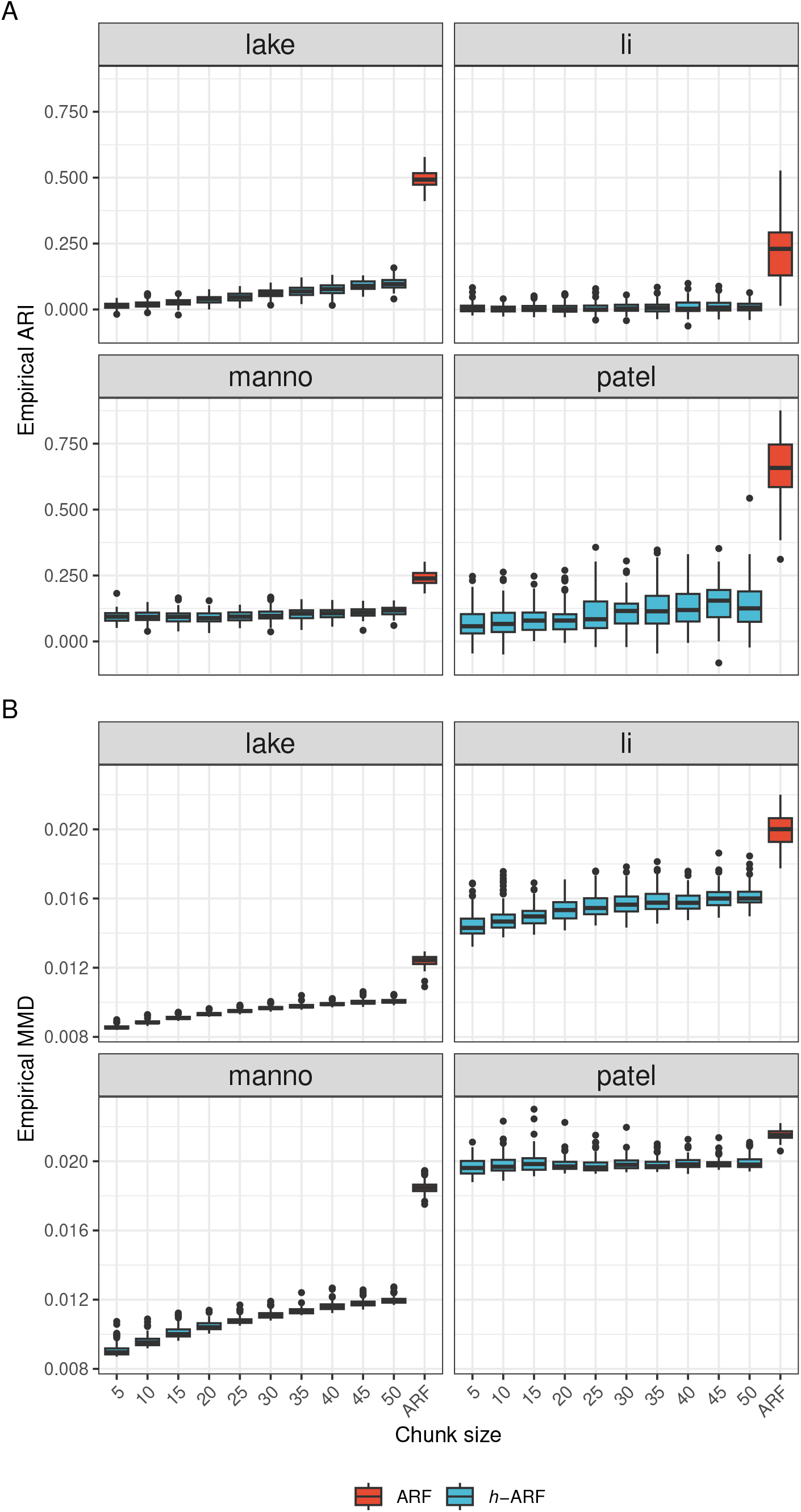
Empirical A) ARI and B) MMD are shown for downstream clustering datasets for different *h*-ARF chunk sizes and for ARF.

**Fig 6.**
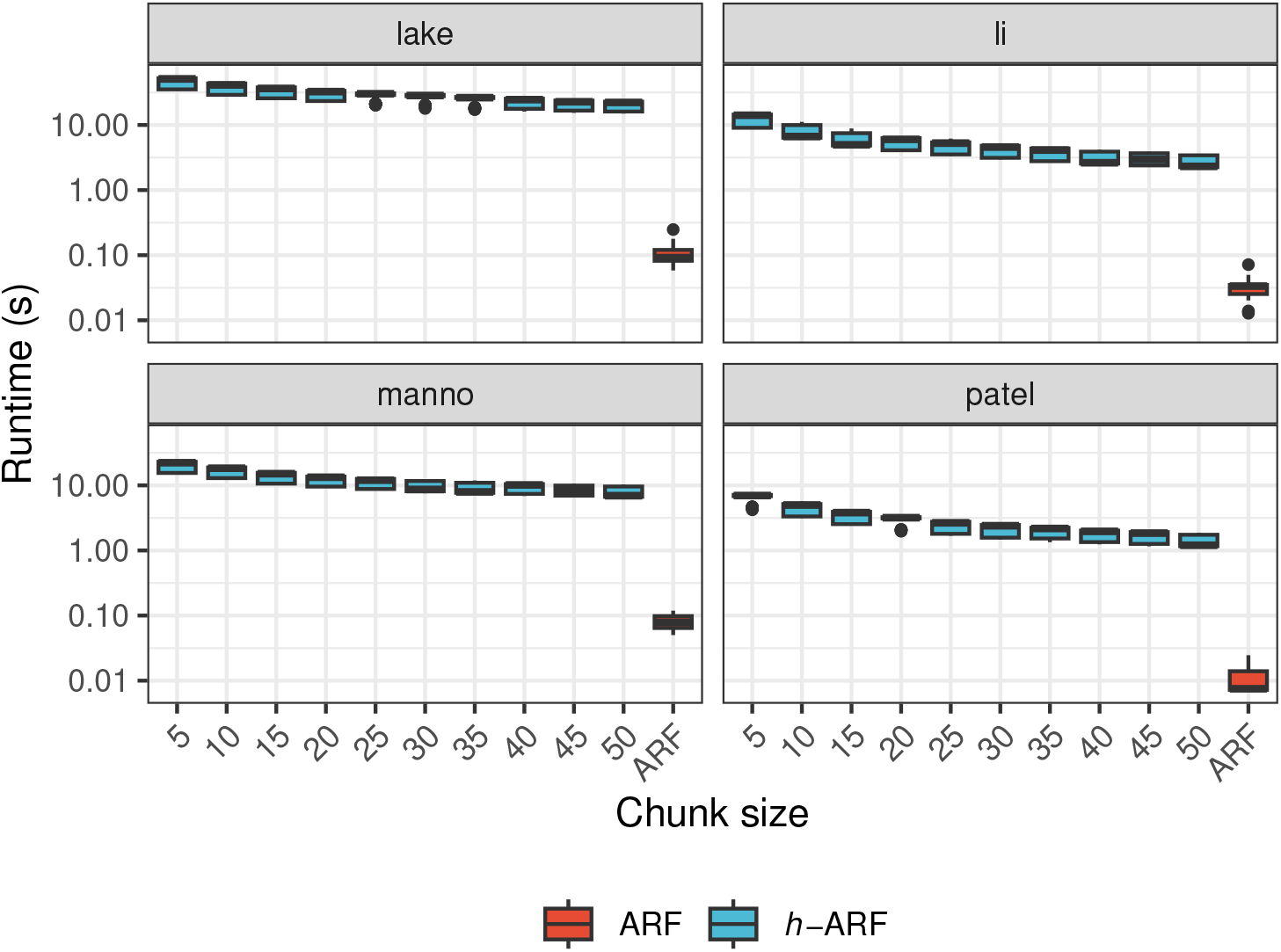
The log10-transformation (for better visualization) of empirical runtimes are shown for downstream clustering datasets for different *h*-ARF chunk sizes and for ARF.

As in the clustering setting, we assess the effect of chunk size as well. Larger chunk sizes reduce distribution fidelity (Figure 7), while having limited impact on predictive performance

**Fig 7.**
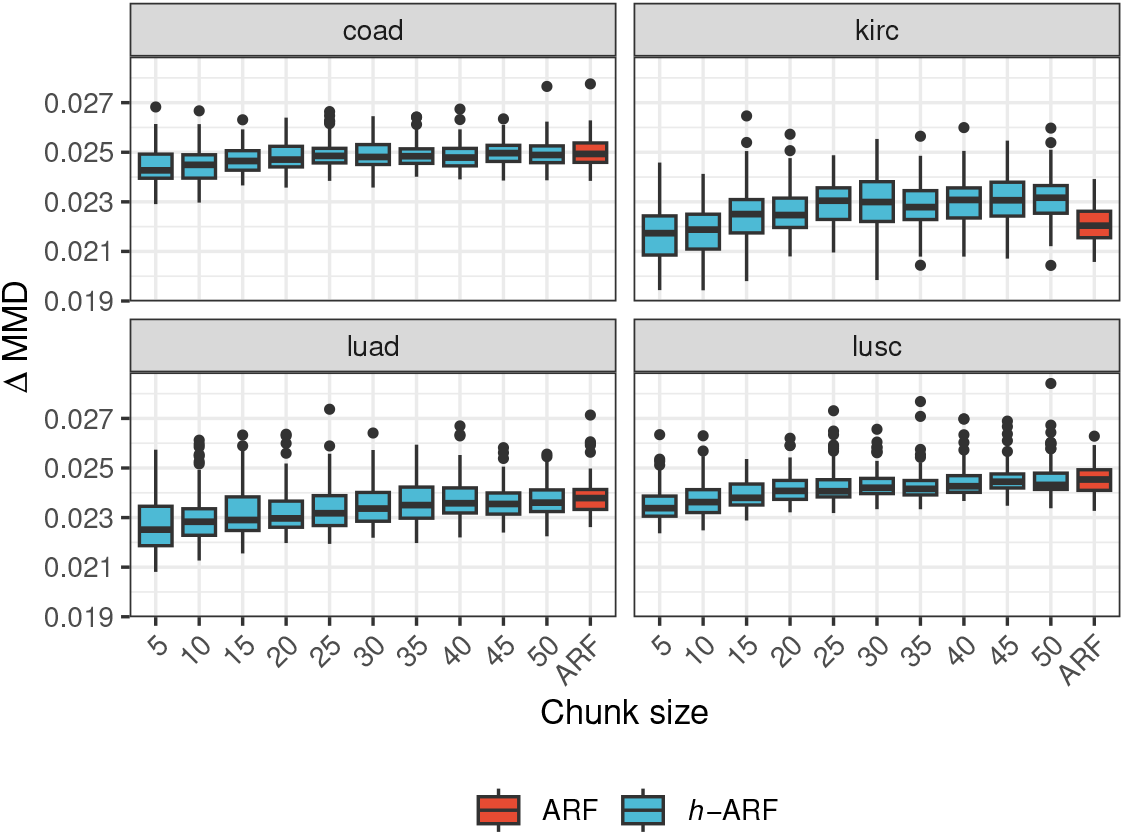
Empirical MMD values are downstream prediction datasets for different *h*-ARF chunk sizes and for ARF.

**Fig 8.**
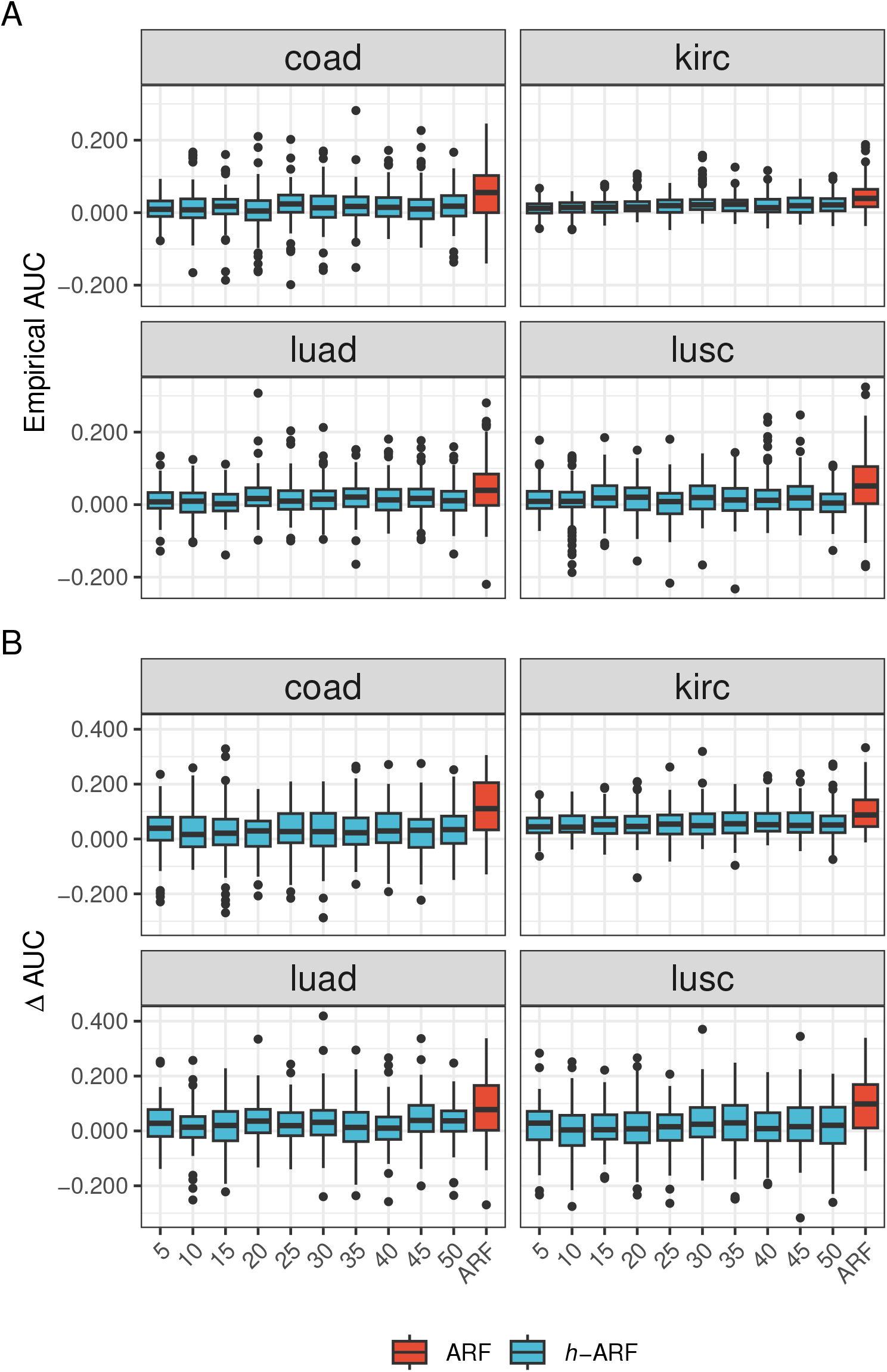
Empirical AUC values are shown for A) RF and B) Lasso classifiers for downstream prediction datasets for different *h*-ARF chunk sizes and for ARF.

**Fig 9.**
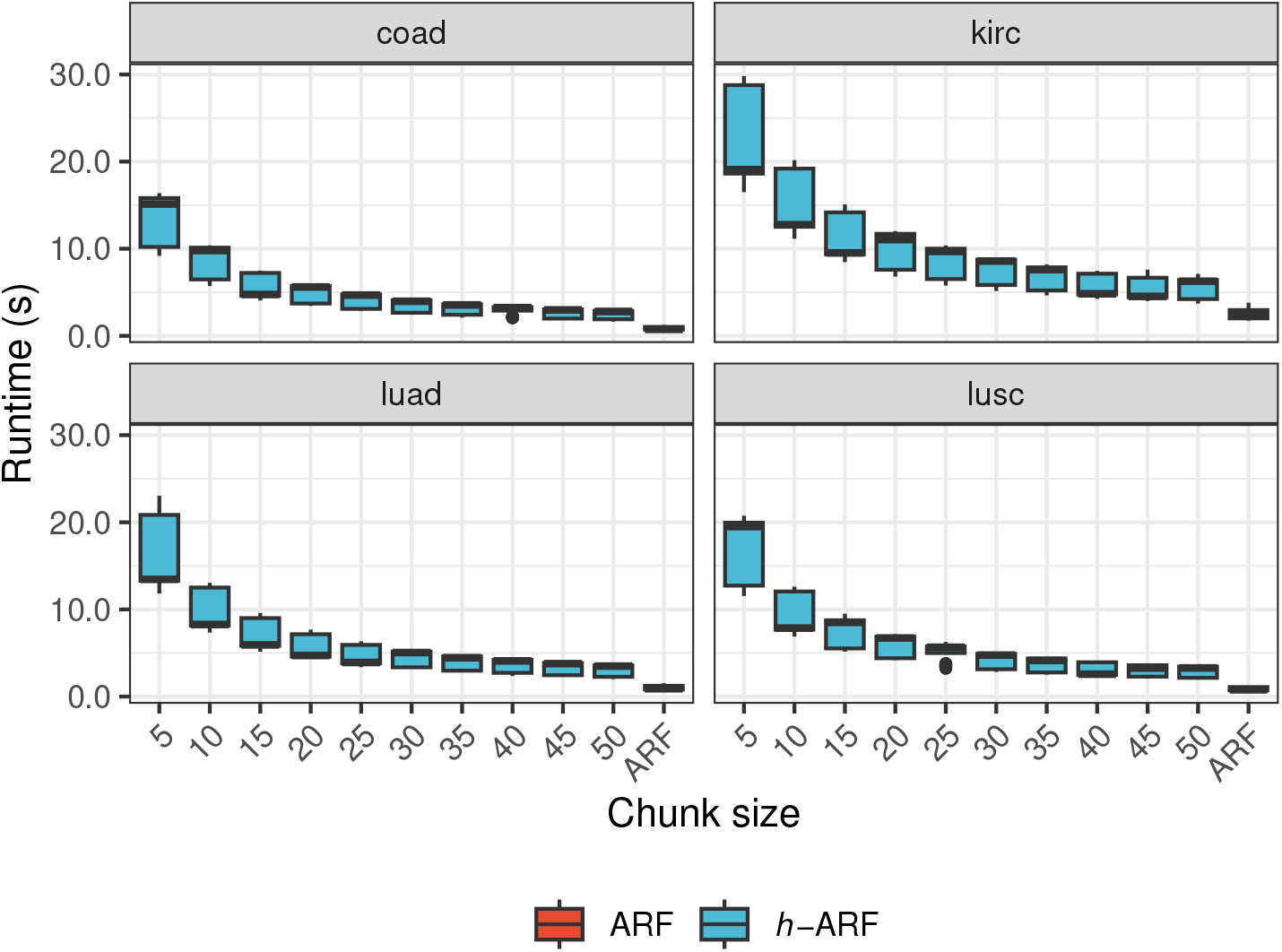
Empirical runtimes for downstream prediction datasets for different *h*-ARF chunk sizes and for ARF.

## 5 Case Study: Alzheimer’s disease brain metabolomics

This section evaluates ARF and *h*-ARF using Alzheimer’s disease (AD) brain metabolomics data. AD is a progressive neurodegenerative disorder and the leading cause of dementia worldwide (45). However, direct molecular profiling and definitive neuropathologic characterization of brain tissue require postmortem samples (46), limiting the availability of brain tissue and, consequently, the sample sizes of brain metabolomics studies. We use data from the Emory University Goizueta AD Research Center brain bank (47), comprising 142 donors with metabolomic measurements and clinical/genetic variables, including sex, age, and *ApoE*. Written informed consent was obtained for all donors. Samples were obtained using research protocols approved by the Emory University Institutional Review Board. These data have previously helped identify metabolomic features and pathways associated with established AD neuropathology markers (48). As response variable for this study, we utilize the Braak’s score (49), ranging from 0 (no pathology) to 3 (severe neocortical involvement), but dichotomized into low (≤ 2) and high (> 2) pathology.

Metabolomic features are treated as the omics modality, while sex, age, and *ApoE* represent clinical covariates. High-resolution metabolic profiling was conducted using liquid chromatography–high-resolution mass spectrometry (LCHRMS), on prefrontal cortex tissue, and with previously established protocols as described by (48). Given the strong association between the *ApoE ε*4 allele and AD risk (50), we include it in our downstream modeling to investigate how well the original effect will be preserved in synthesized data. Metabolic features were filtered using their empirical variance distribution to isolate the top 20% most variable metabolites, reducing high-dimensional sparsity and bringing the total to 7, 205 features. Following this filtering step, the selected metabolites were normalized using training-set feature scaling.

We evaluate ARF and *h*-ARF over 50 runs using a 70:30 train–test split. Both models use 10 trees, with *h*-ARF set to a chunk size of 15. Synthetic data utility is assessed using random forest (5000 trees) and Lasso regression, evaluated via AUC and Matthews Correlation Coefficient (MCC) (51). Lasso models are constrained to avoid shrinkage of the *ApoE* effect. Performance is compared on test data between models trained on original and synthetic data.

Figure 4 reports (A) downstream predictive performance (MCC and AUC), (B) distribution-based similarity metrics (UVD, MMD, and CD), and (C) estimated *ApoE* odds ratios. On the original data, RF and Lasso achieve mean MCCs of 0.40 and 0.46, and AUCs of 0.69 and 0.73, respectively. Overall, *h*-ARF consistently outperforms ARF in downstream predictive utility, with models trained on *h*- ARF synthetic data closely matching those trained on the original data, whereas ARF substantially degrades predictive signal. Distributional metrics show comparable performance between methods, with similar UVD, slightly lower MMD for *h*-ARF, and marginally better CD for ARF. Both methods preserve the direction of the *ApoE ε*4 effect, but only *h*-ARF maintains a realistic effect size, whereas ARF leads to substantial attenuation.

## 6 Conclusion

We introduced the *h*-ARF algorithm as an extension of ARF for high-dimensional generative modeling. We identified key limitations of ARF in high-dimensional settings, including potential non-convergence and violations of local independence assumptions in terminal nodes. To address this, *h*-ARF partitions the feature space into lower-dimensional chunks where ARF assumptions are more likely to hold. Dependencies between chunks are preserved using latent representations, currently obtained via canonical correlation analysis (CCA), which primarily captures linear relationships (27). Future extensions could incorporate more flexible encoding strategies. The parameters of *h*-ARF include chunk size, which could be tuned after assessing the correlation structure of the data. Alternatively to tuning, users could precede the *h*-ARF algorithm with biological network analysis to identify meaningful groups of omics features and inform the choice of the number of clusters, *k*. We recommend the R package `WGCNA` for this purpose (52). Additionally, low-dimensional graphical representations of the omics data can guide the selection of the number of latent components while reducing tuning time. Based on our experience, no more than four latent components were required. While *h*-ARF outperformed ARF on feature distribution–based metrics in the downstream clustering datasets, both methods showed comparable performance in the prediction datasets. We hypothesize that this difference is mainly driven by variations in sample size and signal-to-noise ratio, where signal is defined as the joint dependency structure among features. The runtime is a substantial trade-off of the *h*-ARF framework, as the method requires training ARF models separately within chunks, followed by conditional sampling across partitions.

## 7 Conflicts of interest

The authors declare that they have no competing interests.

## 8 Funding

This work was supported by the German Research Foundation (DFG: #KU 4686/1-1 and # KU 4686/2-1). The brain metabolomics work was supported by the HERCULES Pilot Project via NIEHS P30ES019776 (Anke Hüls), via NIA P30 AG055611 and NIH Grant R01AG087250 (Anke Hüls/Donghai Liang) and the Rollins School of Public Health Dean’s Pilot and Innovation Grant (Anke Hüls).

## 9 Data availability

The data underlying this article are publicly available. The single-cell transcriptomics datasets used in the downstream clustering analyses can be downloaded from the Hemberg Lab GitHub repository: https://github.com/hemberg-lab/scRNA.seq.datasets. Gene expression datasets (Coad, Luad, Lusc, and Kirc) used in the downstream prediction analyses are available through the `curatedTCGAData R` package (version 1.32.1). Requests for the brain metabolomics data should be submitted through the Goizueta Alzheimer’s Disease Research Center at Emory University via the following link: https://alzheimers.emory.edu/research/for-researchers/data-request-form.html.

The code implementing the *h*-ARF algorithm is available on GitHub at https://github.com/bips-hb/harf, and the corresponding R package is available on CRAN. The code required to reproduce all results presented in this article is available at https://github.com/bips-hb/harf-paper.

Operating system: Platform independent Programming language: `R`, version 4.3.0 License: GPL-3 Restrictions for non-academic use: none.

## 10 Author contributions statement

C.J.K.F.: Funding, Conceptualization, Methodology, Software, Writing – original draft. JK.: Conceptualization, Methodology, Software, Validation, Writing–Review & Editing. AH: Conceptualization, Writing–Review, Editing & Data. DL: Writing–Review, Editing & Data. MNW: Conceptualization, Methodology, Software, Validation, Writing–Review.

## 11 Acknowledgments

TCGA data were obtained from the TCGA Research Network, and Alzheimer’s disease brain data from the Emory University Goizueta AD Research Center (ADRC). We gratefully acknowledge the brain donors, research volunteers, and staff of the Goizueta ADRC at Emory University for their participation and contributions.

## Supplementary Material

This document contains the supplementary materials for the article Adversarial Random Forests for Omics Synthesis.

## 12 Benchmark studies

### 12.1 Downstream clustering

Figure 5 shows the impact the empirical ARI and MMD for different values the of chunk sizes in the *h*-ARF algorithm. Because the effect of the chunk size was similar for all performance metric, we show only the empirical MMD for distributional similarity metric and the empirical ARI and downstream analysis utility. In figure 6, we show the runtime of *h*-ARF and ARF for chunk variation.

### 12.2 Downstream prediction

As the tendance was the same for MMC, we only show the empirical AUC values against the chunk size variation (figure 8). Analaguously, we show the impact of chunk size on MMD only, as the behaviour was similar with the other distribution metrics (7).

## Notes

### Competing Interest Statement

The authors have declared no competing interest.

